# Transcriptional profiling of transport mechanisms and regulatory pathways in rat choroid plexus

**DOI:** 10.1101/2022.02.21.481301

**Authors:** Søren N. Andreassen, Trine L. Toft-Bertelsen, Jonathan H. Wardman, Rene Villadsen, Nanna MacAulay

**Author notes:** To whom correspondence should be addressed: Nanna MacAulay, University of Copenhagen, Faculty of Health and Medical Sciences, Department of Neuroscience, Blegdamsvej 3, DK-2200 Copenhagen N, Denmark.

## Abstract

**Background:** Dysregulation of brain fluid homeostasis associates with brain pathologies in which fluid accumulation leads to elevated intracranial pressure. Surgical intervention remains standard care, since specific and efficient pharmacological treatment options are limited for pathologies with disturbed brain fluid homeostasis. Such lack of therapeutic targets originates, in part, from the incomplete map of the molecular mechanisms underlying cerebrospinal fluid (CSF) secretion by the choroid plexus.

**Methods:** The transcriptomic profile of rat choroid plexus was generated by RNA Sequencing (RNAseq) of whole tissue and epithelial cells captured by fluorescence-activated cell sorting (FACS), and compared to proximal tubules. The bioinformatic analysis comprised mapping to reference genome followed by filtering for type, location, and association with alias and protein function. The transporters and associated regulatory modules were arranged in discovery tables according to their transcriptional abundance and tied together in association network analysis.

**Results:** The transcriptomic profile of choroid plexus displays high similarity between sex and species (human, rat, and mouse) and lesser similarity to another secretory epithelium, the proximal tubules. The discovery tables provide lists of transport mechanisms that could participate in CSF secretion and suggest regulatory candidates.

**Conclusions:** With quantification of the transport protein abundance in choroid plexus and their potentially linked regulatory modules, we envision a molecular tool to devise rational hypotheses regarding future delineation of choroidal transport proteins involved in CSF secretion and their regulation. Our vision is to obtain pharmaceutical targets towards modulation of CSF production in pathologies involving disturbed brain water dynamics.

## Introduction

The brain is bathed in cerebrospinal fluid (CSF) that occupies the ventricular system, the subarachnoid space, and the interstitial space between structures and cells in the brain. The CSF serves to create buoyancy for the brain, to protect it from mechanical insult, and as a route by which metabolites, nutrients, and hormones can disperse within the brain [1]. The CSF is produced at a rate of 500 ml per day in adult humans [2], and the majority of the CSF secretion takes place across the choroid plexus [3], which is a specialized secretory tissue located in each of the ventricles. The choroid plexus consists of a monolayer of tight junction-connected epithelial cells, which rest on highly vascularized stroma with connective tissue [4].

A range of cerebral pathologies, i.e. hydrocephalus, stroke and subarachnoid hemorrhage, associate with elevated intracranial pressure (ICP). If left untreated, the brain tissue and the vasculature within compress, further reducing blood flow to the affected areas. Elevated ICP can occur following brain fluid accumulation arising either by reduced drainage of CSF following an obstruction in the brain fluid exit pathways or by hypersecretion of CSF, the latter of which has been observed in conditions such as choroid plexus hyperplasia, choroid plexus papilloma, and in a rodent model of posthemorrhagic hydrocephalus [5–7]. Elevated ICP is routinely treated by insertion of a ventriculo-peritoneal shunt or by a craniectomy [8, 9]. Although these are life-saving procedures, they are highly invasive and associated with severe side effects. Targeted and efficient pharmaceutical treatment aimed at reducing CSF secretion, and thus balancing the brain fluid content, is a desired addition to the clinical toolbox. However, such pharmaceutical approaches have, so far, generally failed due to intolerable side effects or lack of efficiency [10, 11].

Although the existence and production of CSF have been acknowledged for more than a century, the molecular mechanisms underlying this fluid secretion remain unresolved. Some choroidal transport mechanisms have been implicated in the CSF secretion, but their quantitative contribution and the molecular mechanism by which the fluid is transported from the vascular compartment to the brain ventricles await determination [2,3,12]. Importantly, a complete map of the choroidal transport proteins may reveal other fluid-secreting transport mechanisms that could serve as future choroid plexus-specific pharmaceutical targets aimed at reducing CSF secretion in pathological conditions that would benefit from such treatments.

Here we performed transcriptomic analysis of rat choroid plexus from male and female rats and created a searchable database on the obtained transcriptomic profiles. To reveal putative future pharmacological targets, transport mechanisms and regulatory pathways were identified, ranked according to expression levels, and tied together in association networks.

## Materials and Methods

### Experimental rats

This study conformed to the European guidelines and ethical regulations for use of experimental animals. The study utilizes 9-week-old Sprague Dawley rats (Janvier Labs, France) of male and female sex. The rats were housed with 12:12 light cycle with access to water and food ad libitum in accordance with the guidelines of the Danish Veterinary and Food administration (Ministry of Environment and Food) and approved by the animal facility at the Faculty of Health and Medical Sciences, University of Copenhagen. The rats were anaesthetized with intraperitoneal injection of xylazine and ketamine (6 mg/ml and 60 mg/ml in sterile water, 0.17 ml per 100 g body weight (ScanVet, Fredensborg, Denmark)) prior to decapitation and tissue collection.

### Isolation of choroid plexus and proximal tubules

Choroid plexus (from lateral and 4^th^ ventricles) were isolated from five male and five female rats, pooled respectively and stored in RNAlater® (Sigma-Aldrich, St. Louis, Missouri, USA) at -80 °C. Kidney tissue was collected from the male rats, minced, and subsequently digested for 25 min at 37°C in a table shaker at 850 rpm in collagenase solution containing 1 mg/ml collagenase (type II, Gibco®, Grand Island, NY, USA) and 1 mg/ml pronase (Roche, Mannheim, Germany) in buffer solution containing (in mM): 140 NaCl, 0.4 KH_2_PO_4_, 1.6 K_2_HPO_4_, 1 MgSO_4_, 10 Na-acetat, 1 α-ketoglutarat, 1.3 Ca-gluconat, 5 glycin, in addition to 48 mg/l aprotinin (trypsin inhibitor, Sigma-Aldrich, St. Louis, Missouri, USA) and 25 mg/l DNase I (grade II, Roche, Mannheim, Germany), pH 7.56. In five-minute intervals, 1 ml of the solution containing the kidney tissue was transferred to an eppendorf tube containing 1 ml cold buffer solution with 0.5 mg/ml bovine serum albumin (Sigma-Aldrich, St. Louis, Missouri, USA) and replaced by 1 ml collagenase solution. Proximal tubules were manually collected under a microscope, centrifuged at 600×g for 5 minutes, and the pellet stored in RNAlater® at -80 °C.

### Fluorescence-activated cell sorting (FACS) of choroid plexus epithelial cells

Choroid plexus was isolated from 10 male rats, minced, and digested in collagenase (15 mg/ml collagenase (type II, Gibco®, Grand island, NY, USA) in artificial CSF (aCSF)-HEPES containing (in mM): 120 NaCl, 2.5 KCl, 3 CaCl_2_, 1.3 MgSO_4_, 1 NaH_2_PO_4_, 10 glucose, 17 Na-HEPES, pH 7.56 for 30 min at 37°C in a table shaker at 800 rpm. The supernatant was removed after 5 min centrifugation (600×g) and the pelleted cells were resuspended in aCSF-HEPES. The cells were triturated 20 times with a 1000 µl pipette and filtered through a 70 μm filter (pluriStrainer, Mini 70 µm, PluriSelect, Leipzig, Germany), prior to incubation with an anti-NKCC1 antibody with an extracellular epitope (1:200 in aCSF-HEPES, #ANT-071, Alomone Labs™, Jerusalem, Israel) for 30 minutes at 4°C. The cells were pelleted (600×g, 5 min) and resuspended in secondary antibody (1:500 in aCSF-HEPES, Alexa Fluor® 647 - A-21245, Invitrogen™, Carlsbad, California, USA), in which it was kept for 20 min at 4°C prior to centrifugation (600×g, 5 min) and resuspension in cold aCSF-HEPES. Cells were analyzed and sorted on a FACSAria Fusion flow cytometer (BD Biosciences, Lyngby, Denmark).

### Immunohistochemistry

15 μl FACS suspension was placed on poly-D-lysine-coated coverslips for 30 min at room temperature, after which excessive liquid was removed and the attached cells covered with 4% paraformaldehyde in PBS for 15 min at room temperature. Coverslips were washed 3 times with 0.02% tween-20 in PBS (PBST) and permeabilized with PBST for 10 min at room temperature. Cells were treated with a blocking solution (4 % normal goat serum (NGS) in PBST) for 1 hour at 4°C prior to exposure to primary antibody against AQP1 (1:400, #AQP-001, Alomone Labs™, Jerusalem, Israel) at 4°C O/N. Coverslips were washed with PBST and incubated with secondary antibody (1:700, A-11034, Alexa Fluor® 488, Invitrogen™, Carlsbad, California, USA) and Phalloidin (1:400, A22287, Alexa Fluor™ 647, Invitrogen™, Carlsbad, California, USA) for 2 hours at room temperature. Cells were washed and mounted onto microscope glass slides with ProLong™ Gold Antifade Mountant with DAPI (P36935, Invitrogen™, Carlsbad, California, USA).

### RNA extraction and sequencing

The RNA extraction and library preparation were performed by Novogene Company Limited, UK with NEB Next® Ultra™ RNA Library Prep Kit (NEB, USA) prior to their RNA sequencing (paired-end 150 bp, with 12 Gb output) on an Illumina NovaSeq 6000 (Illumina, USA).

### Bioinformatics and computational analyses

All program parameter settings for library building and mapping, together with all scripts for the gene annotation and analysis are available at https://github.com/Sorennorge/MacAulayLab-RNAseq2. All analyses exclude genes transcribed at levels below a cut-off at 0.5 transcripts per million (TPM) [13]. Raw data are avaliable at the National Center for Biotechnology Information (NCBI) Gene Expression Omnibus (GEO) database (accession number: GSE194236).

### RNA sequencing analysis

The 150 base paired-end reads were mapped to reference genome (Rattus norvegicus Rnor_6.0 v.103) using Spliced Transcripts Alignment to a Reference (STAR) RNA-seq aligner (v. 2.7.2a) [14]. The mapped alignment by STAR was normalized to TPM with RSEM (RNA-Seq by Expectation Maximization v. 1.3.3) [15]. Gene information was gathered with mygene (v3.1.0) python library [16–18], from which gene symbol, alias, and Gene Ontology (GO) terms [19–21] were extracted. Mitochondrial genes (seqname: MT) were omitted from the reference genome prior to comparison of the kidney proximal tubule transcriptional profiles with that of the choroid plexus, since the greater abundance of these genes in the proximal tubule would skew the plasma membrane transporter expression comparison.

### Cross-species comparison

The human choroid plexus transcriptome was obtained from GEO database (GSE137619, SRR10134643-SRR10134648) [22–24] and the mouse choroid plexus transcriptome was obtained from GEO (GSE66312, SRR1819706-SRR18197014) [25]. All samples were quality controlled with fastqc [26] and trimmed with Trimmomatic [27] (Slidingwindow 4:20, minimum length of 35 bp). The human and mouse samples, together with rat sample 3 (male), were mapped to the human reference genome (Homo sapiens GRCh38 v.104), the mouse reference genome (Mus musculus GRCm39 v.104), and rat reference genome (Rattus norvegicus Rnor_6.0 v.103) with STAR (v. 2.7.2a). The reference genome for rat, mouse, and human for the cross-species analysis was only containing gene/transcripts of biotype ‘protein coding’. Mapped alignments were normalized to TPM with RSEM (v. 1.3.3.) and the mean from the samples for human and mouse, respectively, were used for further analysis. Transcribed genes sharing gene name between the compared species (or their orthologue, collected from ensemble.org via biomart martview) were included in the cross-species comparison.

### Category section

Gene lists of transporters, pumps, water and ion channels, and G protein-coupled receptors (GPCR) were collected from the ‘target and family list’ from Guide to Pharmacology [28–31]. Genes annotated as ‘transporters’ were employed to generate the list of membrane transporters and pumps whereas genes annotated as ‘voltage-gated ion channels’, ‘ligand-gated ion channels’, and ‘other ion channels’ were employed to generate the list of water and ion channels. To filter for plasma membrane proteins, the transporter and pump gene list was initially filtered to exclude the mitochondrial and vacuolar transport families SLC25, ATP5, and ATP6V, after which the transporter and channel lists were filtered based on associated GO terms; ‘integral component of plasma membrane’ or ‘plasma membrane’, but only included genes annotated as ‘integral component of membrane’ or ‘transmembrane’, but not annotated as ‘lysosome’, ‘endosome membrane’, ‘lysosomal’, ‘mitochondrion’, ‘mitochondrial’, ‘golgi apparatus’, ‘vacuolar’, or ‘endoplasmic’. Genes annotated as ‘GPCR’ were employed to generate the list containing GPCRs. Receptor tyrosine kinases (RTK) were gathered from Human Genome Organisation (HUGO) Gene Nomenclature Committee (HGNC) database [32], with the annotations ‘receptor tyrosine kinase’ including sub group ‘ephrin receptors’ and ‘ErbB family’. The list of kinases was obtained from the Kyoto Encyclopedia of Genes and Genomes (KEGG) database [33–36] for entries of ‘EC 2.7.10.2’ (non-specific protein-tyrosine kinase), ‘EC 2.7.12’ (Dual-specificity kinases) with the two sub-categories, and ‘EC 2.7.11’ (Protein-serine/threonine kinases) with the 33 sub-categories. These three entries were collected with organism specific “rno” (Rattus norvegicus) filter. These kinases were filtered for protein kinases by GO terms: ‘protein kinase activity’, ‘protein serine/threonine kinase activity’, ‘protein serine kinase activity’, ‘protein threonine kinase activity’, ‘protein tyrosine kinase activity’, ‘map kinase activity’. Kinases involved solely with transcription or cell cycle modulation were subsequently excluded. Phosphatases were gathered from the KEGG database with entries EC 3.1.3 (Phosphoric Monoester Hydrolases) with 108 subcategories. The phosphatases were filtered for protein-interacting phosphatases by GO terms: ‘phosphoprotein phosphatase activity’, ‘protein serine/threonine phosphatase activity’, ‘protein serine phosphatase activity’, ‘protein threonine phosphatase activity’, and ‘protein tyrosine phosphatase activity’. Phosphodiesterases (PDE) were collected based on gene name starting with ‘PDE’. Cyclases were obtained from the KEGG database [33–36] entry numbers ‘EC 4.6.1.1’ (adenylate cyclase) and ‘EC 4.6.1.2’ (guanylate cyclase).

### Network analysis

The network analysis was generated from protein-protein association tables from the String-database [37, 38] as a plugin for Cytoscape (v. 3.8.2) [39]. Firstly, full interaction tables were generated through a full protein query of every protein in the lists of transporters and pumps, water and ion channels, GPCRs, RTKs, kinases, phosphatases, PDEs and cyclases. Secondly, the tables were filtered for interaction between the ‘transporters and pumps’ and all the regulatory genes (lists of GPCRs, RTKs, kinases, phosphatases, PDEs, and cyclases). All interactions between regulatory proteins were discarded. The same was done for ‘water and ion channels’. We only included protein-protein associations that were curated in a database or were demonstrated to interact experimentally. Thirdly, these interaction tables were loaded back into Cytoscape (v. 3.8.2) and modulated: Confidence score (from string-db) of 0.6-1 was used for genes from the list ‘transporter and pumps’ interactions and 0.7-1 was used for genes from the list ‘water and ion channels’.

## Results

To characterize the choroid plexus ‘transportome’, we obtained the transcriptomic profile of choroid plexus from adult male and female Sprague-Dawley rats by performing RNAseq of the tissue. The data are organized in a searchable webserver-based database (https://cprnaseq.in.ku.dk/) to allow search for genes of interest. The database encompasses the genes expressed in the choroid plexus, their expression level in TPM, and their alias (protein name, to the extent that this feature was available).

### The transcriptomic profile of rat choroid plexus mimics those of mouse and human

To determine the species similarity of the choroidal transcriptomic profile, we compared that obtained from the rat to published versions of these obtained from human [22] and murine [25] choroid plexus. The rat shares 89% of its choroidal protein-coding genes with that of the human choroid plexus, and 91% with that of mice (Fig. 1). These numbers may represent an underestimate, as some of the remaining genes might not be annotated with the shared name or as an orthologue, or may fall just above or below the cut-off level of 0.5 TPM. However, the 11% non-shared expressed genes in humans only account for 3.3% of the total transcripts and only 5.5% of these non-shared genes were annotated with ‘transport activity’, meaning that only 0.1% are transporters not shared with the rat. The 9% non-shared expressed genes for mouse account for 2.8% of the total transcripts, and only 0.1% was annotated as transporters. Such transcriptomic similarity suggests that the functional integrity is conserved among these organisms, and that the rat is an applicable animal model in which to determine physiological aspects, especially those that are membrane transport-related, of the choroid plexus.

**Fig 1.**
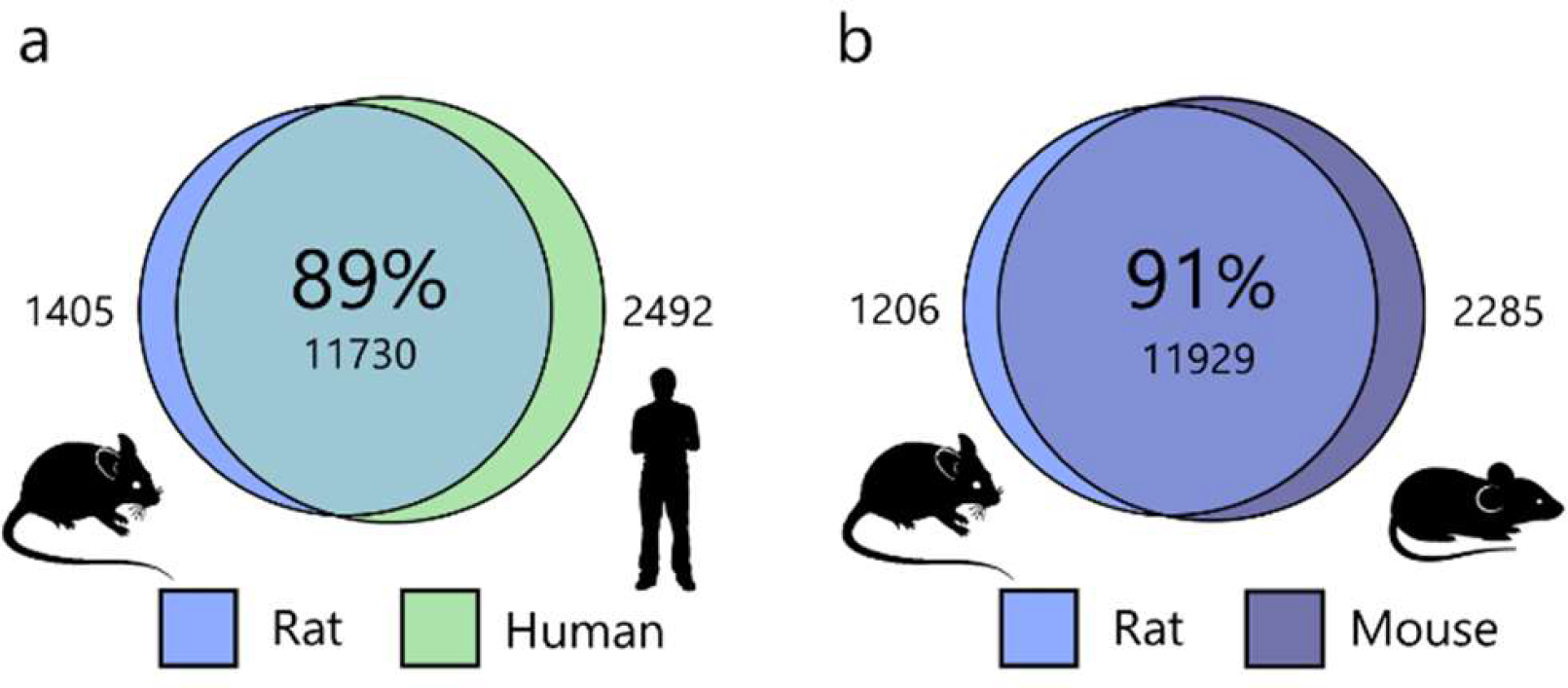
Species comparison of transcribed genes in choroid plexus. RNAseq analysis reveals common protein coding transcribed genes of rat choroid plexus compared with a human [22] and b mouse [25]. Transcribed genes are included as shared if they share the same protein name or are listed as orthologue genes. The percentage in the sphere center refers to rat perspective, with non-shared genes on each side of the overlapping spheres.

### Transport proteins highly expressed in choroid plexus may be involved in CSF secretion

To obtain a complete list of transport proteins expressed in the choroid plexus and their transcriptomic abundance, the RNAseq data were filtered for genes encoding transport proteins located in the plasma membrane (see Methods). Each gene was associated with its alias, and a description of the type of membrane transport protein. Some protein names and their function were well established, while other transport mechanisms expressed in choroid plexus appeared not fully characterized with no general agreement on a given alias or function. The lists were therefore manually curated according to the Universal Protein Resource (Uniprot) [40] and the Human Gene Database (Genecards) [41] to ensure that the widest accepted alias and function were associated with each gene name. The transport proteins are divided into i) coupled transporters and ATP-driven pumps and ii) water and ion channels, and sorted according to their choroidal transcript abundance (Supplementary Table 1 and 2). Table 1 and 2 illustrate the 20 highest expressers in each category. Several of the transport proteins implicated in CSF secretion are found amongst these highly expressed transport proteins; different subunits of the Na^+^/K^+^-ATPase, the Na^+^,K^+^,2Cl^-^ cotransporter (NKCC1), aquaporin 1 (AQP1), and the HCO_3_^-^ transporters NBCe2, NCBE, and AE2 [12, 42]. In addition, the lists present some additional transport proteins that could be envisaged to partake in CSF secretion, i.e. cation, anion and sulfate transporters (BOCT, BSAT1, SNAT1, SUT1, and MCT8, Table 1) in addition to various K^+^ and Cl^-^ ion channels (Table 2) with no prior association to CSF secretion. With such high expression in this tissue, these transport proteins are likely to serve a physiologically important function in choroid plexus, and could potentially be involved in CSF secretion.

**Table 1.**
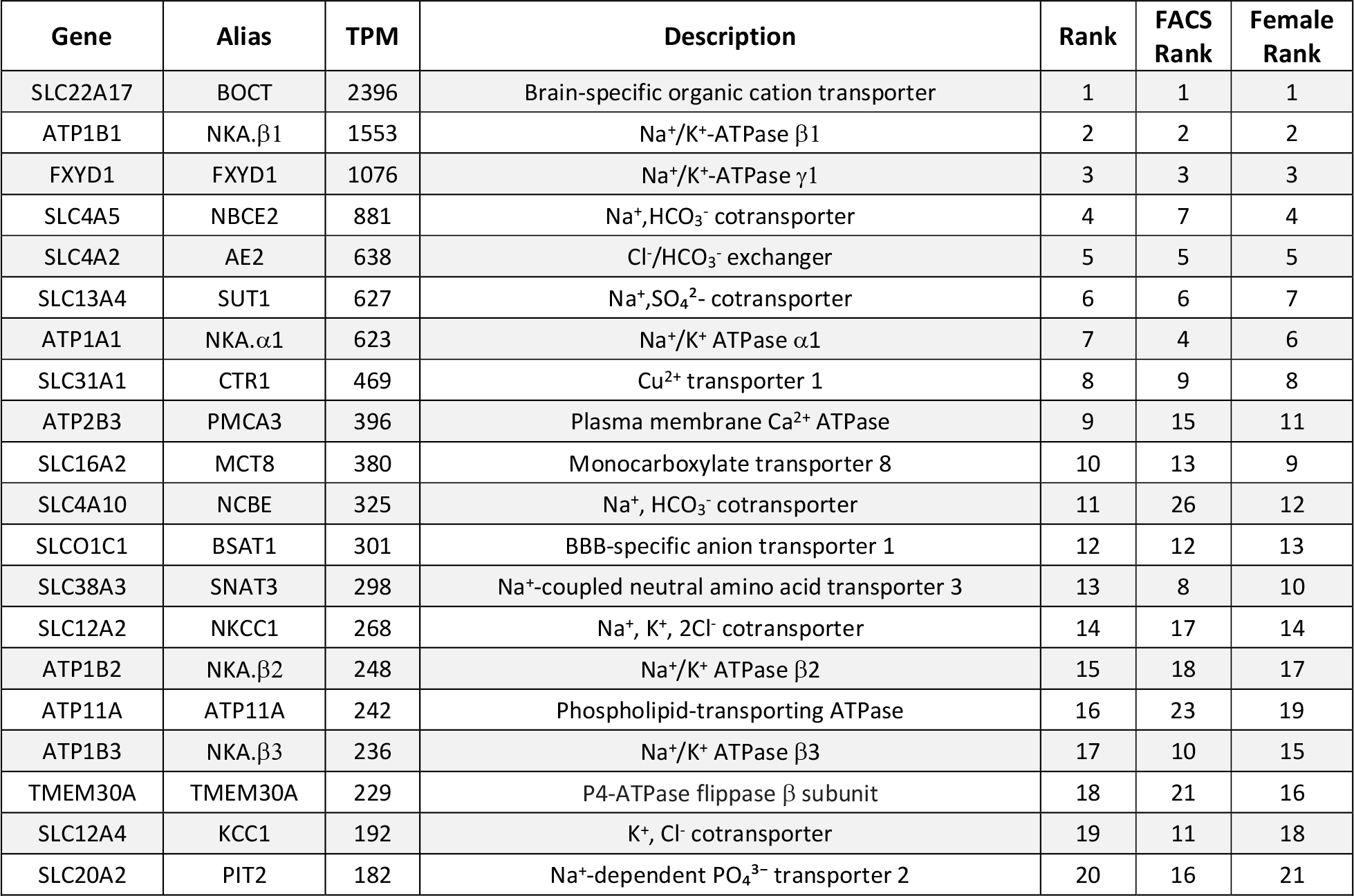
Highly transcribed transporters and pumps in choroid plexus. RNAseq analysis revealing the 20 highest expressed genes encoding plasma membrane transporters and pumps in choroid plexus. TPM; transcripts per million, FACS rank; each gene’s rank in the FACS sample, Female rank; rank in the choroid plexus from female rats.

**Table 2.**
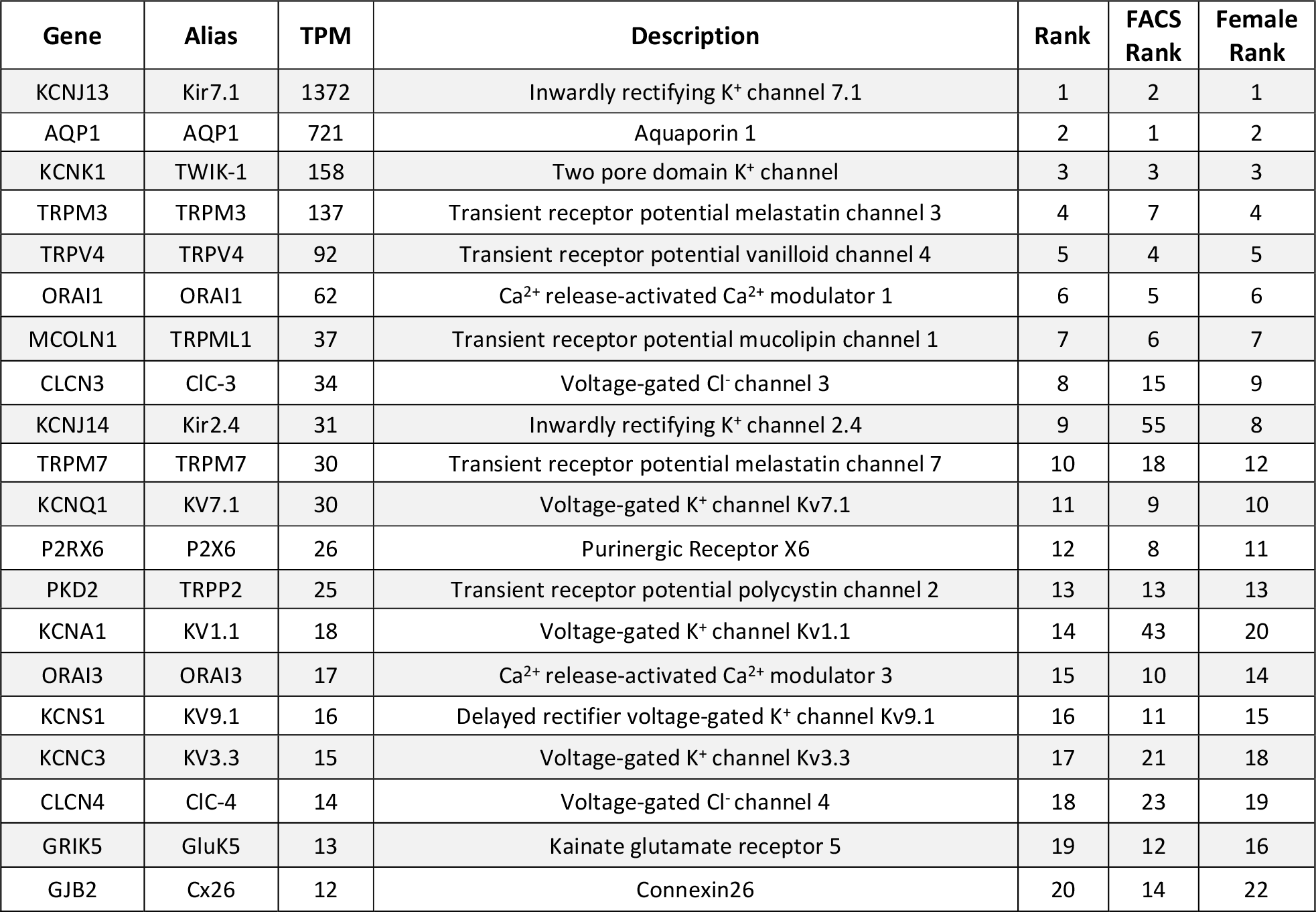
Highly transcribed membrane channels in choroid plexus. RNAseq analysis revealing the 20 highest expressed genes encoding plasma membrane channels in choroid plexus. TPM; transcripts per million, FACS rank; each gene’s rank in the FACS sample, Female rank; rank in the choroid plexus from female rats.

### The transcriptome of a pure fraction of choroid plexus epithelial cells is comparable to that of choroid plexus

The choroid plexus is a feather-like structure containing a monolayer of epithelial cells with centrally located vasculature, stroma, and immune cells. Most RNAseq studies of this tissue take advantage of the fact that the majority (around 90%) of the cells in the tissue are choroid plexus epithelial cells [43, 44] and thus include the entire structure into the RNAseq procedure [45–49]. In this manner, some of the transcripts are anticipated to arise from other cell types than that of the choroid plexus epithelium. To resolve a putative discrepancy between a pure fraction of choroid plexus epithelial cells and that of the entire tissue, dissociated cells from acutely isolated choroid plexus were captured by FACS. FACS with the anti-NKCC1 antibody targeted to the extracellular face of this choroidal transport protein produced a fraction of highly fluorescent and large cells (Fig. 2A), which was absent in the FACS conducted in the absence of antibody (Fig. 2B) or with only the secondary antibody (Fig. 2C). Intact choroid plexus epithelial cells captured by FACS expressed AQP1 (Fig. 2D), which in the choroid plexus is expressed solely in the epithelial cells [50], demonstrating their epithelial origin. RNAseq analysis of the epithelial cells captured by FACS retrieved 97% of the genes detected in the entire choroid plexus structure (Fig. 2E). The 3% of the genes that were absent from the choroid plexus epithelial cells captured by FACS, were largely (>95% of the genes with cell type annotation) annotated primarily to cell types other than epithelial cells [51, 52], such as cells of mesenchymal, endothelial, neuronal, immune, or glial origin, which reside in the choroid plexus structure [44]. The cell population obtained from the FACS procedure thus essentially represents choroid plexus epithelial cells. Choroid plexus epithelial cells captured by FACS displayed nearly identical transcriptomic profiles to that of the entire choroid plexus of choroidal transport proteins (97%, Fig. 2F) and channels (93%, Fig. 2G). The non-shared genes place around the cut-off TPM of 0.5 for both transporters and channels. More specifically, the transporters/pumps found amongst the top 20 highest expressers in the choroid plexus were all detected within the top 26 highest expressed transport proteins in the pure fraction of choroid plexus epithelial cells (Table 1). Such similarity was reproduced for choroidal membrane channels (except for the two ion channels K_ir_2.4 and K_v_1.1 that placed further down the list), Table 2. Complete lists of transporters and membrane channels detected in the purified choroid plexus epithelial cells are included as Supplementary Table S3 and S4. We conclude that RNAseq analysis of the complete choroid plexus structure provides a representative quantitative identification of the membrane transport proteins expressed in the choroid plexus epithelial cells.

**Fig 2.**
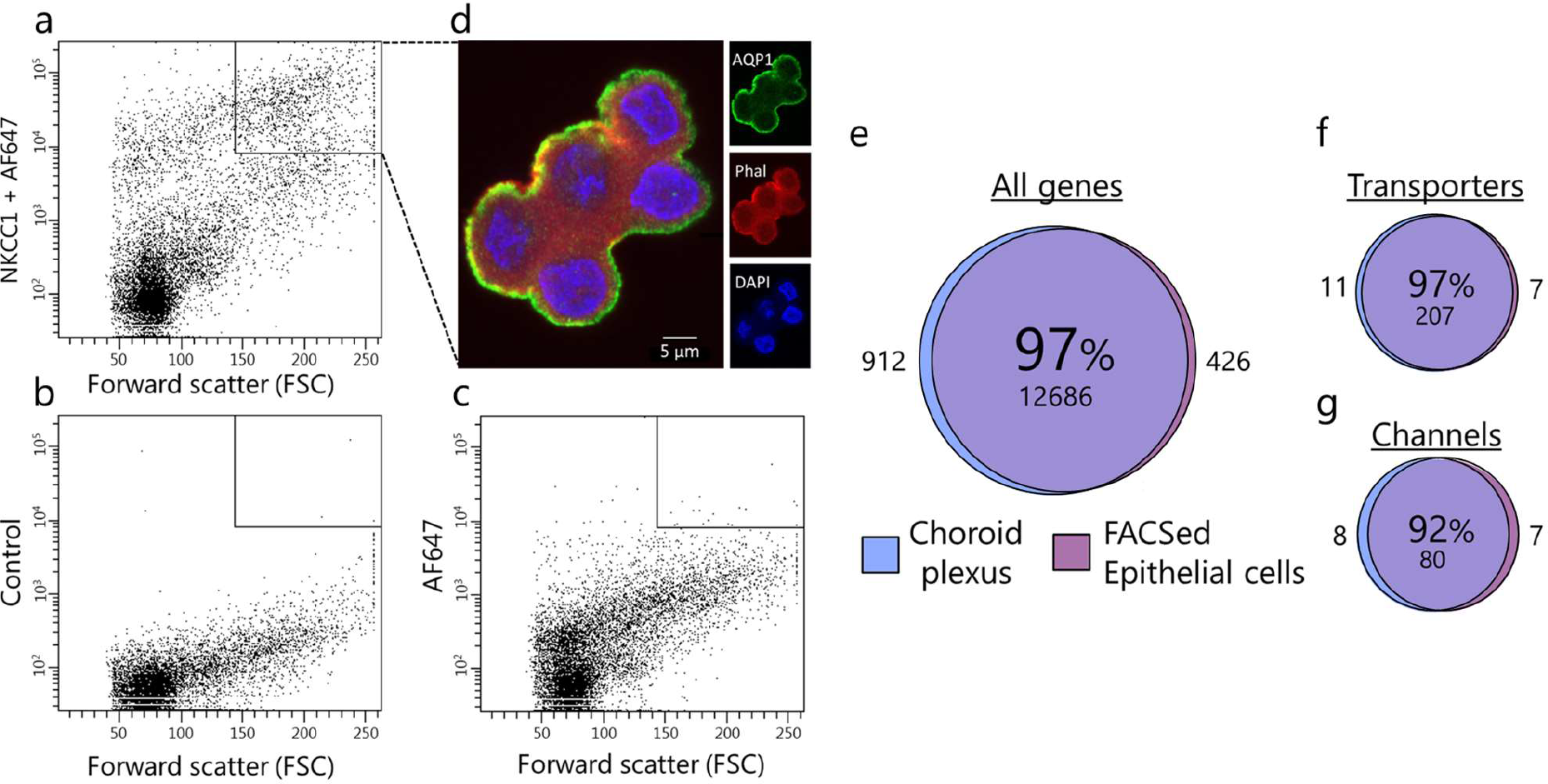
Choroid plexus transcriptomic profile similar to FACS choroid plexus epithelial cells. a-d FACS with fluorescence emission intensity (y-axis) and forward scatter (FSC) (x-axis). a The cell suspension obtained by FACS with primary anti-NKCC1 antibody and secondary antibody, b in the absence of antibody inclusion in the FACS procedure, or c in the absence of primary antibody, but with inclusion of secondary antibody. The target area of capture (cells with high fluorescence emission intensity and large cell size) is defined in a black square in the top right corner (a-c). d Immunohistochemical staining of the captured cells with anti-AQP1 (green), marker of F-actin; phalloidin (red), and nucleus marker; DAPI (blue). e-g Depiction of the shared transcribed genes in the pure fraction of choroid plexus epithelial cells versus the whole choroid plexus, e for all genes, f, for genes encoding membrane transporters and pumps, and g for genes encoding membrane water and ion channels. Shared genes (in percentages and number) in the sphere center, with the number of non-shared genes depicted on either side.

### The choroid plexus expresses a wide range of membrane transporters

The solute carriers (SLC) represent a large group of membrane transport proteins containing more than 400 members divided amongst 66 families [53]. The 66 family names and transport functions were collected from the Bioparadigms SLC database [54, 55]. Of the 66 existing SLC families, transcripts of 52 of these gene families were detected in choroid plexus, and 44 of these coding for SLC family members residing in the plasma membrane (representing 63% of all choroidal transporter and pump transcripts in the plasma membrane). To obtain an overview of choroidal plasma membrane transport functions, the 44 SLC families detected in the choroidal plasma membrane were grouped into 11 supercategories according to similarities in transported substrates (Fig. 3, Supplementary Table S5). The most abundantly expressed supercategory consisted of 31 electrolyte and bicarbonate transporters (six families, 2602 TPM) followed by seven organic an/cationic transporters (one family, 2581 TPM) and 19 large anion transporters (five families, 1929 TPM). These are followed by amino acid and neurotransmitter transporters (36 genes), various sugar transporters (17 genes), and vitamin transporters (6 genes), (Fig. 3 and Supplementary Table S5). Notably, the choroid plexus expresses a wealth of metal transporters (dispersed among 22 genes from five families, 928 TPM, with zinc transporters as the most prominent (15 genes)), (Fig. 3 and Supplementary Table 5). With such abundance and variety in transporter types, the choroid plexus obviously serves important physiological roles in transporting solutes across this epithelial layer, besides that serving as the CSF secreting machinery.

**Fig 3.**
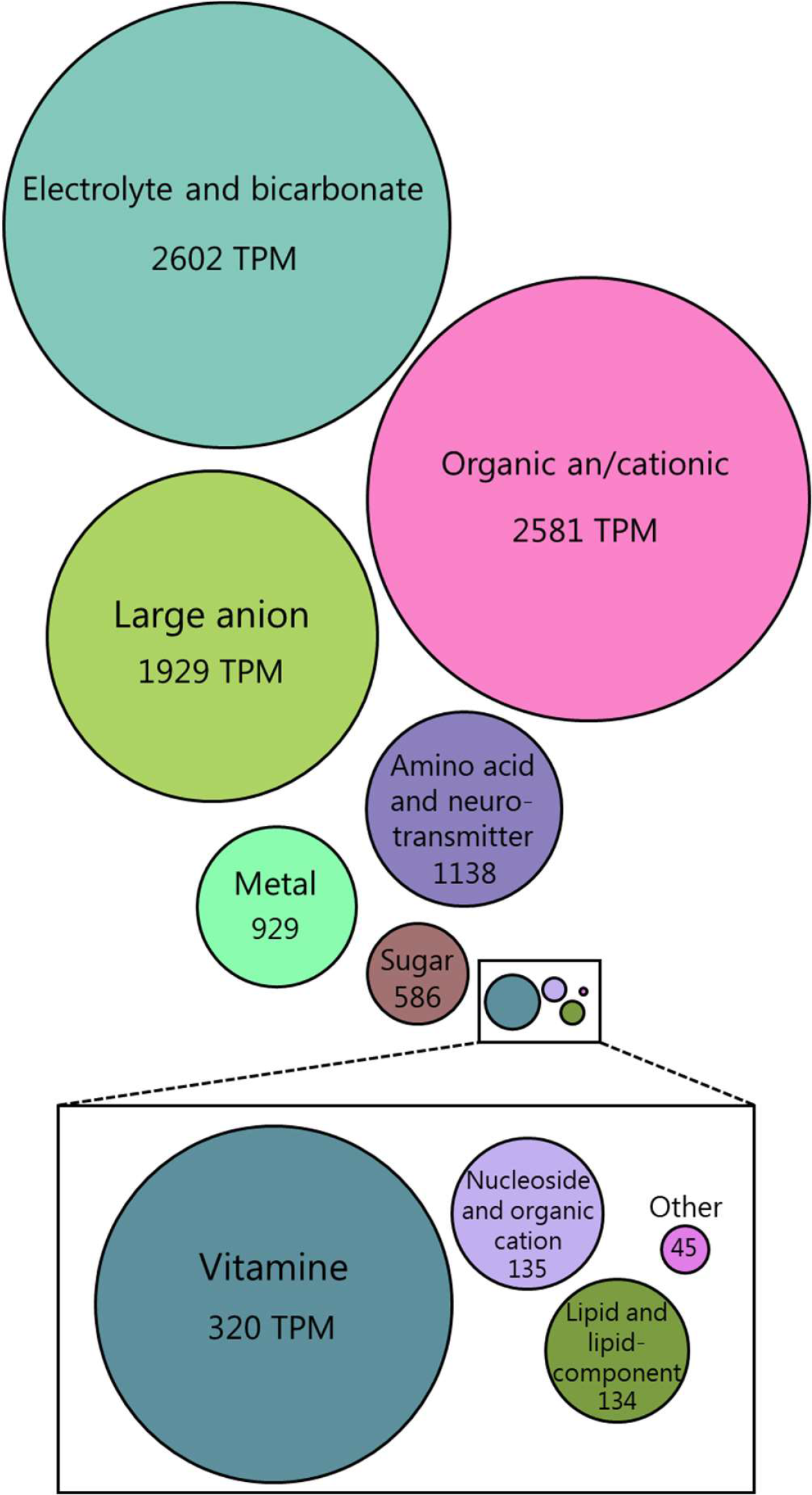
Quantitative expression of genes encoding SLCs. Illustration of the relative transcriptional abundance of the 11 SLC super-categories generated based on similarities in transported substrates (see Supplementary Table S5). The area of each circle illustrates the relative accumulative transcription abundance in TPM (transcripts per million).

### Female and male rats share choroid plexus transcriptomic profile

To verify the sex-specific transcriptomic profile of female versus male rat choroid plexus, we compared the expression profiles for choroid plexus obtained from each age-matched sex. The female choroid plexus shared 98% of the overall expression profile with the male choroid plexus (Fig. 4A), suggesting high functional similarity of the choroid plexus in the two sexes. Notably, the non-shared genes account for less than 0.1% of the total transcripts. To reveal potential sex-specific differences within the CSF secreting machinery, we compared the expression profile for membrane transport mechanisms. Choroid plexus obtained from the female rats shared 98% of the transcripts encoding transporters (Fig. 4B) and 95% of those encoding channels (Fig. 4C) with the male rat. The male 20 highest expressed genes were included in the top 21 highest expressed transporter genes in the female transporter category (Table 1) and in the top 22 highest expressed channel genes in the female channel category (Table 2). Choroid plexus obtained from the two sexes thus shares high degree of expression profile similarity, with near-identical expression of abundant transport proteins. These data suggest a comparable choroidal CSF secreting machinery in female and male rats.

**Fig 4.**
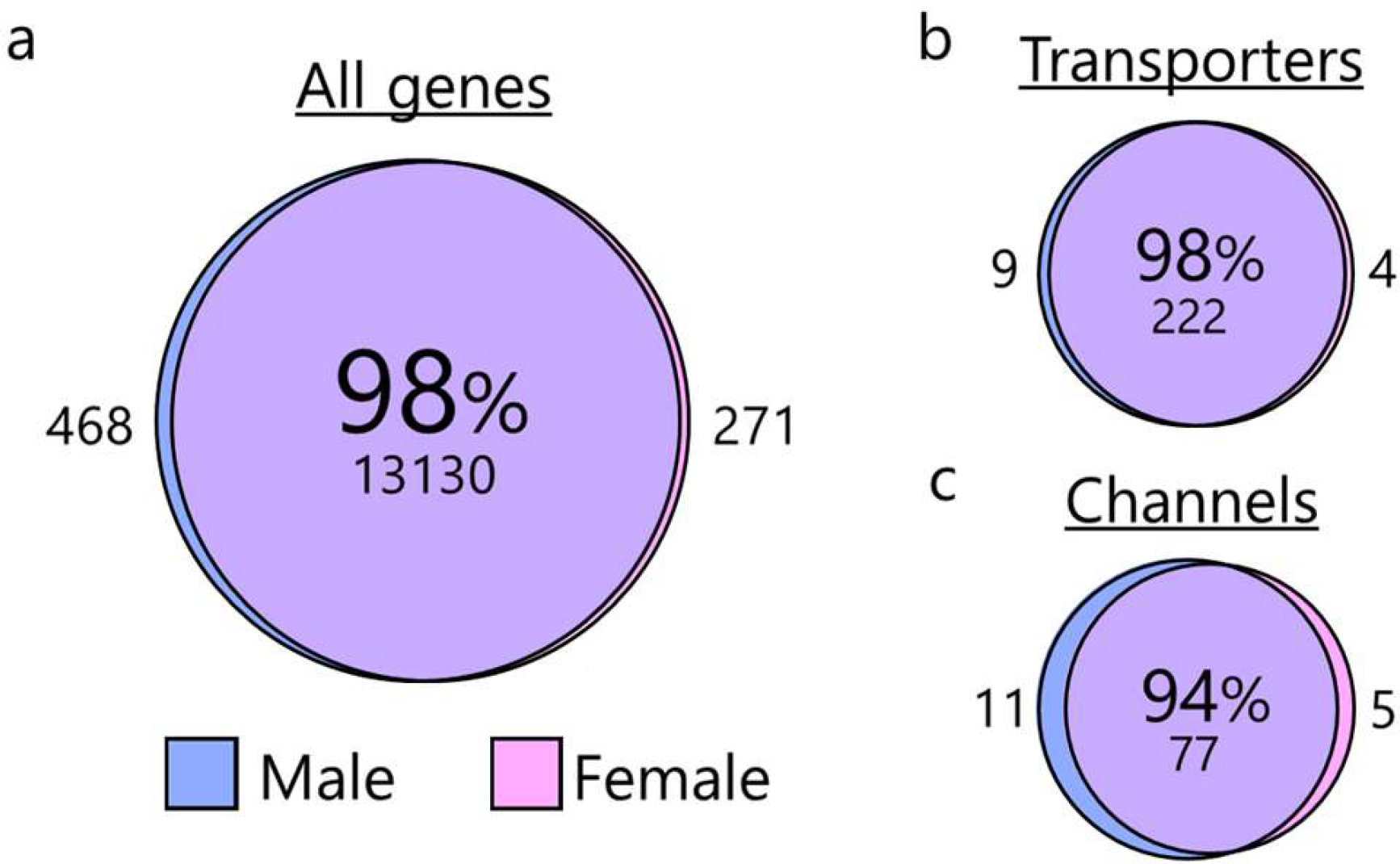
Sex comparison of transcribed genes in choroid plexus. Depiction of the shared transcribed genes in the female choroid plexus versus the male, a for all genes, b for genes encoding membrane transporters and pumps, and c for genes encoding membrane water and ion channels. Shared genes (in percentages and number) in the sphere center, with the number of non-shared genes depicted on either side.

### The choroid plexus transcriptome differs from that of another high-capacity secretory epithelium

To reveal possible transport protein candidates involved in fluid secretion across the choroid plexus, we obtained the transcriptomic profile of an epithelium of comparable secretory capacity to that of choroid plexus; the kidney proximal tubule [2]. The majority (∼91 %) of the transcribed genes obtained in the proximal tubule were detected in the choroid plexus. However with the larger number of transcribed genes found in the choroid plexus, this common pool of transcribed genes amounted to only 41% of the choroidal expression profile (Fig. 5A). 64% of the transporter-encoding transcribed genes obtained from the proximal tubule were retrieved in the choroidal samples (in which proximal tubule transcribed genes represented 30%, Fig. 5B). In the channel category, the equivalent numbers were 52 % of the proximal tubule transcribed genes being retrieved in the choroid plexus samples and 15% of the choroidal transcribed genes detected in the proximal tubule (Fig. 5C). Of the 20 highest expressed transporters/pumps (Table 3) and channels (Table 4) in the proximal tubule (see Supplementary Tables S6 and S7 for the complete lists), only 10 (transporters and pumps) and 10 (water and ion channels) genes were found in the choroid plexus (Tables 3 and 4). The Na^+^/K^+^-ATPase α1β1, AQP1, TRP channels, and some voltage-gated Cl^-^ channels present themselves as common transport mechanisms in the two tissues. Although the proximal tubule joins the choroid plexus in the ranks of highly secretory epithelia, the molecular machinery driving the fluid secretion appears, at least in part, to rely on distinct transport mechanisms.

**Fig 5.**
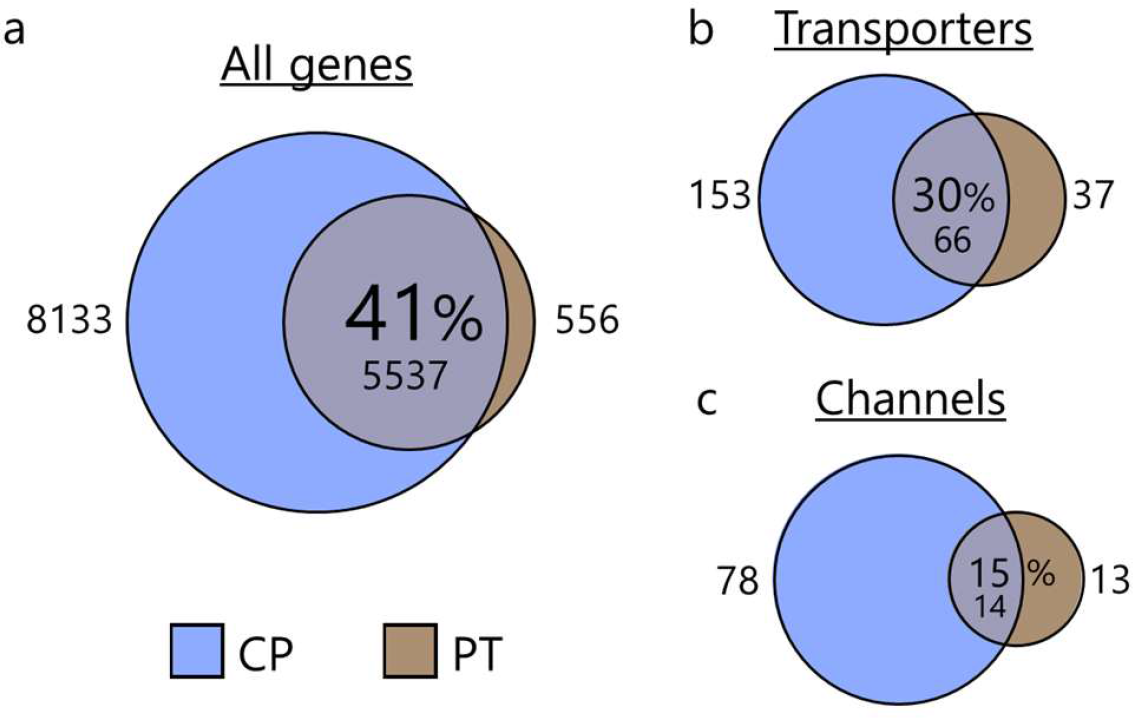
Tissue comparison of transcribed genes in choroid plexus. Depiction of the shared transcribed genes in the choroid plexus (CP) versus those detected in the proximal tubule (PT), a for all genes, b for genes encoding membrane transporters and pumps, and c for genes encoding membrane water and ion channels. Shared genes (in percentages and number) in the sphere center, with the number of non-shared genes depicted on either side.

**Table 3.**
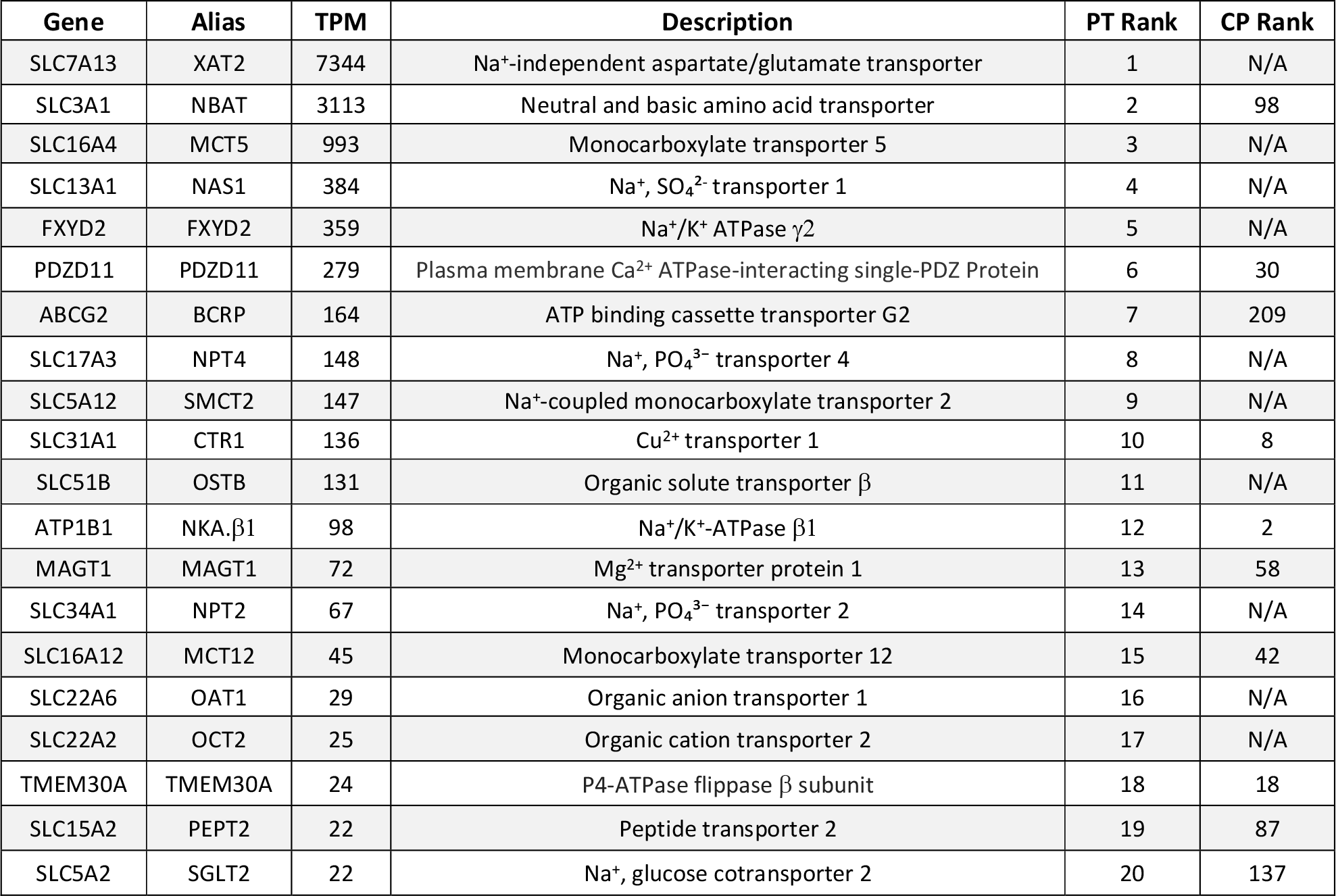
Highly transcribed transporters and pumps in proximal tubule. RNAseq analysis revealing the 20 highest expressed genes encoding plasma membrane transporters and pumps in proximal tubule. TPM; transcripts per million, PT rank; each gene’s rank in the proximal tubule sample, CP rank; each gene’s rank in the choroid plexus sample, N/A; not applicable if the genes is transcribed below the applied cut-off of 0.5 TPM.

**Table 4.**
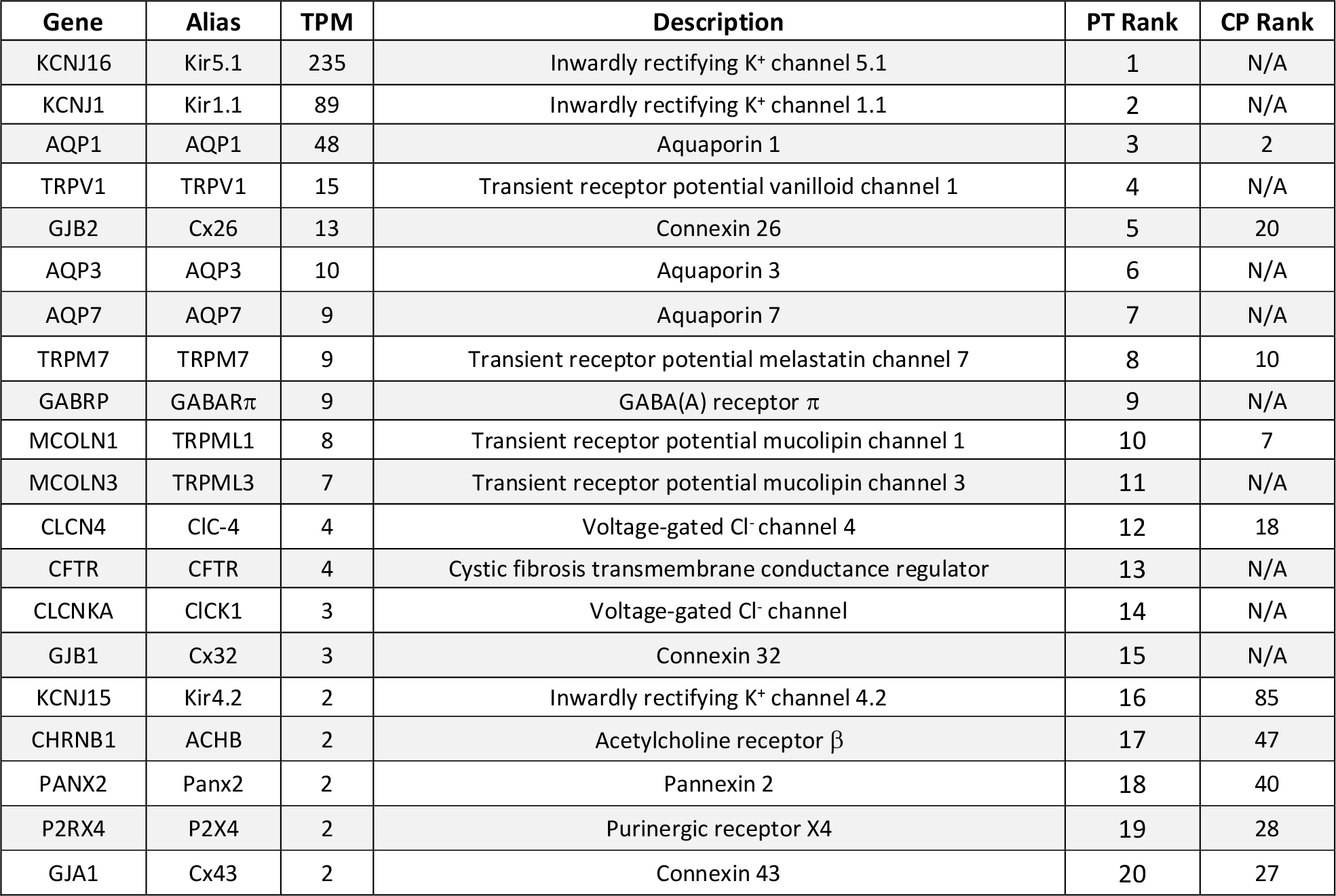
Highly transcribed membrane channels in choroid plexus. RNAseq analysis revealing the 20 highest expressed genes encoding plasma membrane channels in proximal tubule. TPM; transcripts per million, PT rank; each gene’s rank in the proximal tubule sample, CP rank; each gene’s rank in the choroid plexus sample, N/A; not applicable if the genes is transcribed below the applied cut-off of 0.5 TPM.

### Receptors and intracellular modulators may be involved in regulation of CSF secretion

It is anticipated that CSF secretion is tightly regulated to ensure stable brain fluid dynamics and thus intracranial pressure, although the regulatory control of CSF secretion is largely unresolved. To reveal which plasma membrane receptors are expressed in choroid plexus, all G protein-coupled receptors (GPCRs) and receptor tyrosine kinases (RTKs) expressed in choroid plexus were filtered from the RNAseq data set and listed according to their expression levels (Supplementary Tables S8 and S9). Each gene was associated with its alias and a description of the type of receptor (if known). All receptors were manually curated [40, 41] to ensure that the widest accepted alias and function were associated with each gene name. Of the 20 highest expressed GPCRs (Table 5), the serotonin receptor 2C (HTR2C) figures at the top, and is accompanied by the endothelin receptor type B, the GABA_B_ receptor, corticotrophin releasing hormone receptor 2, several adhesion-associated receptors, and those belonging to the family of frizzled GPRCRs. Notably, the list contains several orphan GPCRs (GPR146, GPR175, GPR107) without well-established ligands and functions, although GPR146 has been proposed as a receptor for insulin C-peptide [56], which in turn may regulate the Na^+^/K^+-^ATPase [57]. Dominant amongst the receptor tyrosine kinases top expressers (Table 5) are receptors for growth factors (GF) of various kinds; fibroblast GF, platelet derived GF, vascular endothelial GF, and insulin/insulin-like receptors, in addition to various immune-related RTKs.

**Table 5.**
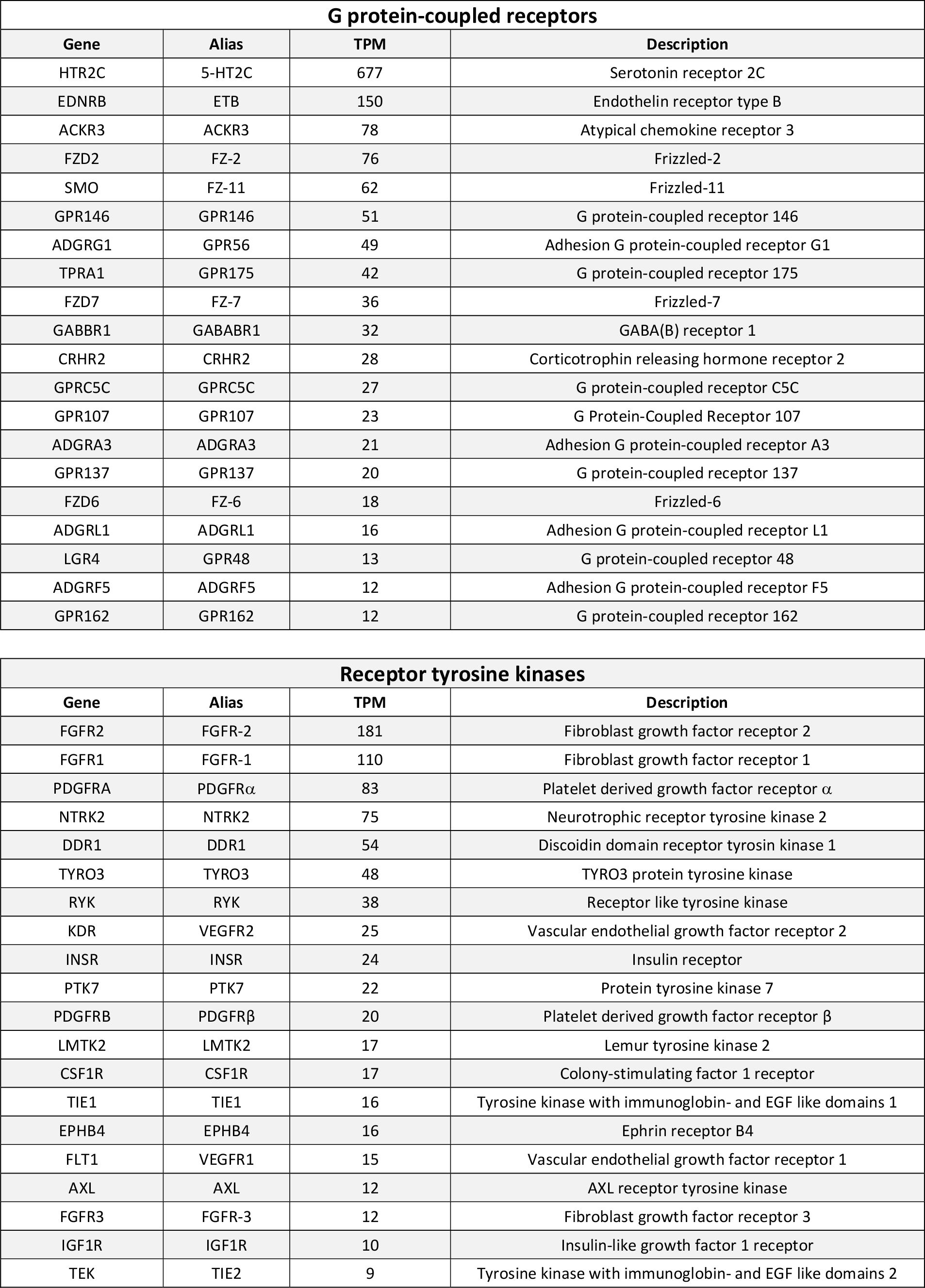
Highly transcribed plasma membrane receptors in choroid plexus. RNAseq analysis revealing the 20 highest expressed genes encoding plasma membrane GPCRs (top) and receptor tyrosin kinases (bottom) in choroid plexus. TPM; transcripts per million.

The receptors generally exert their function via intracellular signaling cascades promoting phosphorylation (kinases) or dephosphorylation (phosphatases) of target proteins such as transport proteins and transcriptional factors potentially promoting expression of select transport proteins. To reveal such regulatory proteins expressed in choroid plexus, the RNAseq data was filtered for kinases and phosphatases, and the data manually curated to obtain alias and function (complete lists found as Supplementary Tables S10 and S11). The list of the 20 highest expressed kinases (Table 6) encompasses a large variety of kinases including MAP kinases, AKT kinases, and casein kinases. Interestingly, the Stk39 kinase (SPAK), which is the second highest expressed kinase in choroid plexus, is directly implicated in CSF secretion via its ability to activate NKCC1 [6]. The list of the 20 highly expressed phosphatases covers a variety of phosphatases encompassing both serine/threonine and tyrosine phosphatases (Table 6).

**Table 6.**
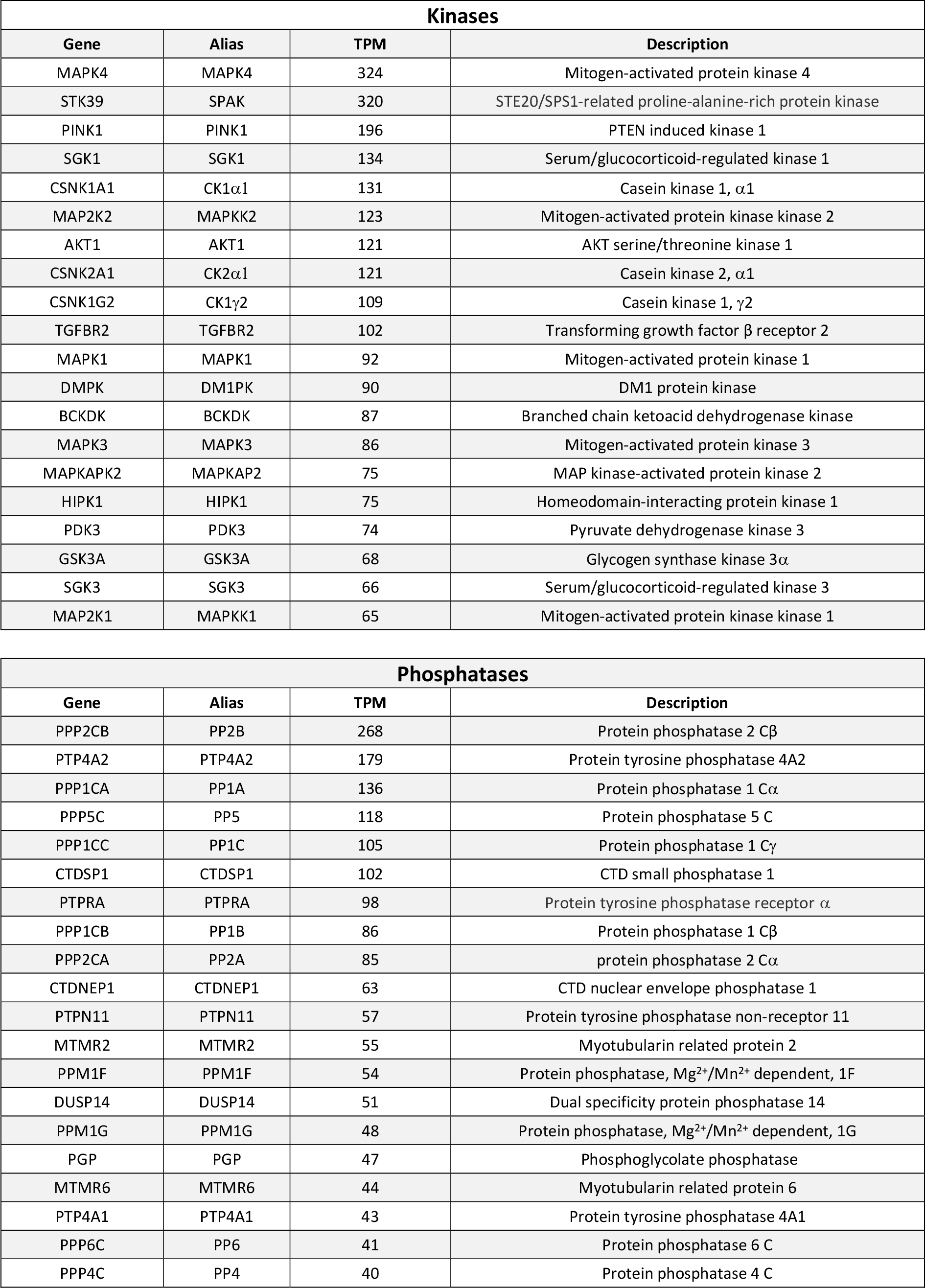
Highly transcribed kinases and phosphatases in choroid plexus. RNAseq analysis revealing the 20 highest expressed genes encoding protein kinases (top) and phosphatases (bottom) in choroid plexus. TPM; transcripts per million.

Some kinases are activated by cyclic nucleotides, such as cAMP required for PKA activation and cGMP required for PKG activation. The abundance of these cyclic nucleotides is regulated by cyclases and phosphodiesterases, the presence and activity of which could well modulate the CSF secretion in the choroid plexus [58–60]. The RNA data set was therefore filtered for the presence of these, and the 10 highest expressers amongst cyclases and phosphodiesterases are listed in Table 7, with the complete lists of manually curated (for alias and function) genes included as Supplementary Tables S12 and S13.

**Table 7.**
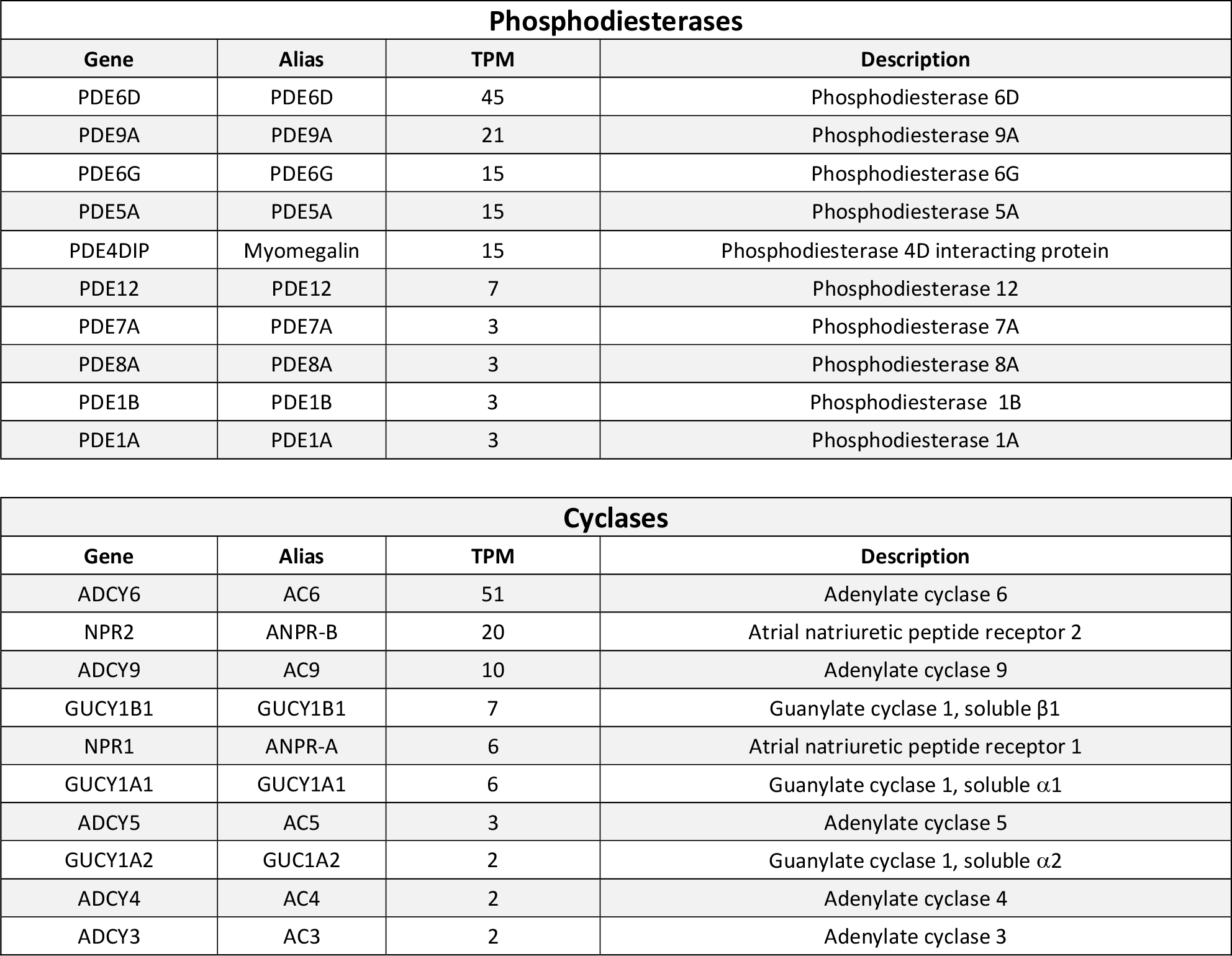
Highly transcribed phosphodiesterases and cyclases choroid plexus. RNAseq analysis revealing the 10 highest expressed genes encoding phosphodiesterases (PDE, top) and cyclases (bottom) in choroid plexus. TPM; transcripts per million.

| Phosphodiesterases |  |  |  |
| --- | --- | --- | --- |
| Gene | Alias | TPM | Description |
| PDE6D | PDE6D | 45 | Phosphodiesterase 6D |
| PDE9A | PDE9A | 21 | Phosphodiesterase 9A |
| PDE6G | PDE6G | 15 | Phosphodiesterase 6G |
| PDE5A | PDE5A | 15 | Phosphodiesterase 5A |
| PDE4DIP | Myomegalin | 15 | Phosphodiesterase 4D interacting protein |
| PDE12 | PDE12 | 7 | Phosphodiesterase 12 |
| PDE7A | PDE7A | 3 | Phosphodiesterase 7A |
| PDE8A | PDE8A | 3 | Phosphodiesterase 8A |
| PDE1B | PDE1B | 3 | Phosphodiesterase 1B |
| PDE1A | PDE1A | 3 | Phosphodiesterase 1A |

| Cyclases |  |  |  |
| --- | --- | --- | --- |
| Gene | Alias | TPM | Description |
| ADCY6 | AC6 | 51 | Adenylate cyclase 6 |
| NPR2 | ANPR-B | 20 | Atrial natriuretic peptide receptor 2 |
| ADCY9 | AC9 | 10 | Adenylate cyclase 9 |
| GUCY1B1 | GUCY1B1 | 7 | Guanylate cyclase 1, soluble β1 |
| NPR1 | ANPR-A | 6 | Atrial natriuretic peptide receptor 1 |
| GUCY1A1 | GUCY1A1 | 6 | Guanylate cyclase 1, soluble α1 |
| ADCY5 | AC5 | 3 | Adenylate cyclase 5 |
| GUCY1A2 | GUC1A2 | 2 | Guanylate cyclase 1, soluble α2 |
| ADCY4 | AC4 | 2 | Adenylate cyclase 4 |
| ADCY3 | AC3 | 2 | Adenylate cyclase 3 |

### Network analysis of choroid plexus

To obtain insight into potential physiologically-relevant regulatory properties of the choroidal transport machinery, we built association networks of transporters/pumps (Fig. 6) and channels (Fig. 7) with the various choroidal receptors (GPCRs and RTKs) and intracellular messengers (kinases, phosphatases, PDEs, and cyclases). Such networks are obtained with a string database and provide links between the transport mechanisms and a regulatory factor, if such has been implied experimentally or in curated databases [61] in published work on any cell type or tissue. Most notably, various isoforms of protein kinase A (PKA) and MAP kinases (MAPK) associate with phosphorylation of different subunits of the Na^+^/K^+^-ATPase and the plasma membrane Ca^2+^-ATPase (PMCA), whereas the facilitative glucose transporter (GLUT4) and the Na^+^/H^+^ exchanger (NHE1) associate with a variety of different regulatory candidates (Fig. 6). The SPAK-mediated regulation of NKCC1 in choroid plexus [6, 62] is evidenced in the network association of these proteins along with the with-no-lysine kinases (WNK1-4), oxidative stress responsive kinase 1 (OSR1), and other cation-Cl^-^ cotransporters (Fig. 6). The different PKA isoforms associate with various ion channels as well, most prominently with voltage- gated K^+^ channels, while PKC isoforms associate with the transient receptor potential vanilloid channel 4 (TRPV4) (Fig. 7), which modulates the rate of CSF secretion [63] and has been implicated in hydrocephalus development in a rat model of genetically-induced hydrocephalus [64]. In addition, various highly expressed phosphatases associate with different ligand- and voltage-gated ion channels (Fig. 7). Altogether, such network analysis indicated a plethora of potential regulatory pathways, some of which could be implicated in regulation of CSF secretion, or other physiological processes.

**Fig 6.**
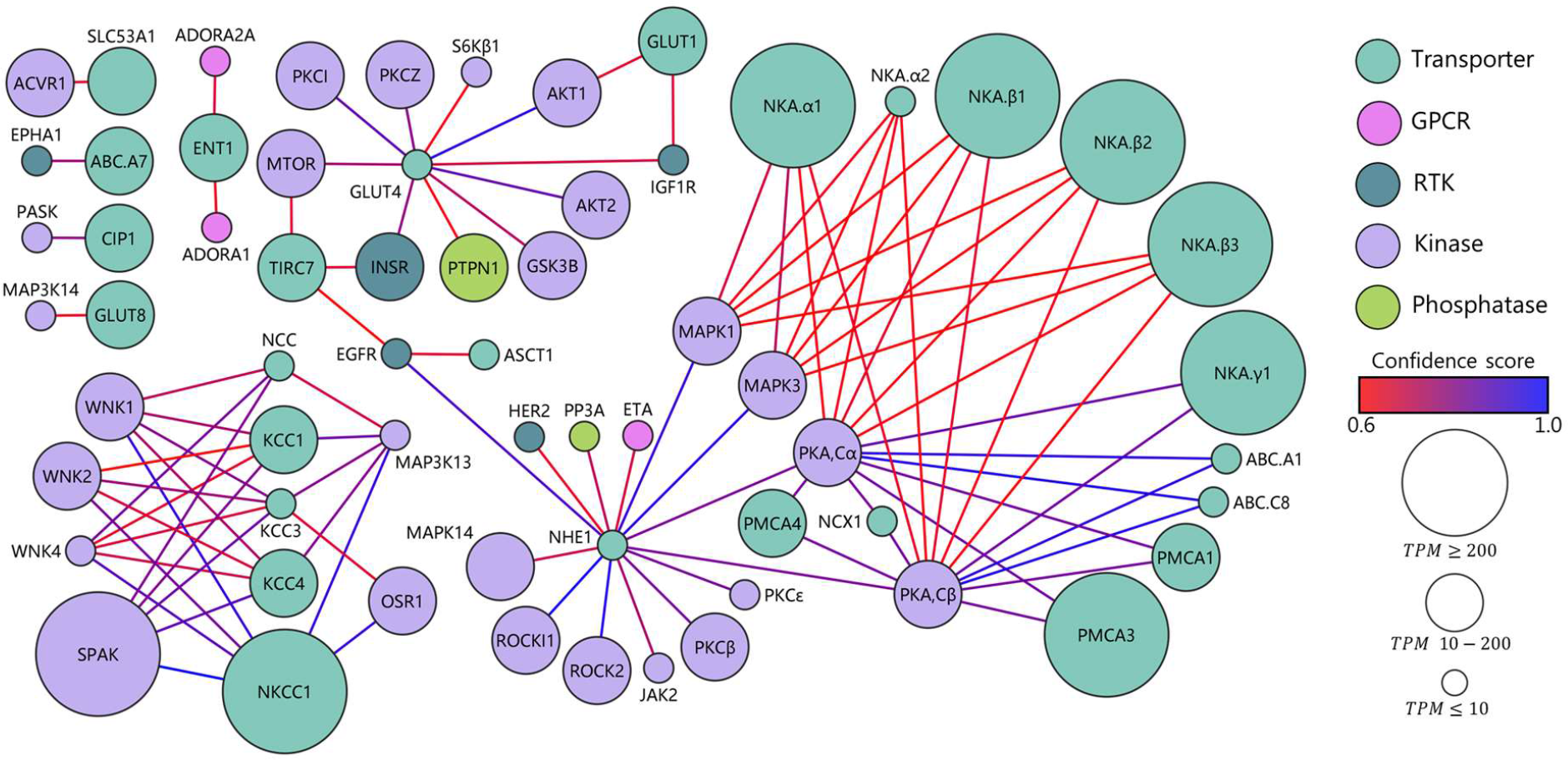
Network analysis of choroidal transporters and signaling cascades. Association network between plasma membrane transporters and pumps together with receptors (GPCRs and RTKs) and intracellular messengers (kinases, phosphatases, PDEs, and cyclases). The nodes (spheres) are colored based on their protein type. The size of the nodes corresponds to the transcriptional expression level in TPM and illustrates TPM ≤ 10, 10 -200, and ≥ 200. The confidence score is indicated by the color of the connecting lines from 0.6 (red) to 1.0 (blue). Only associations from empirical data and curated databases are included.

**Fig 7.**
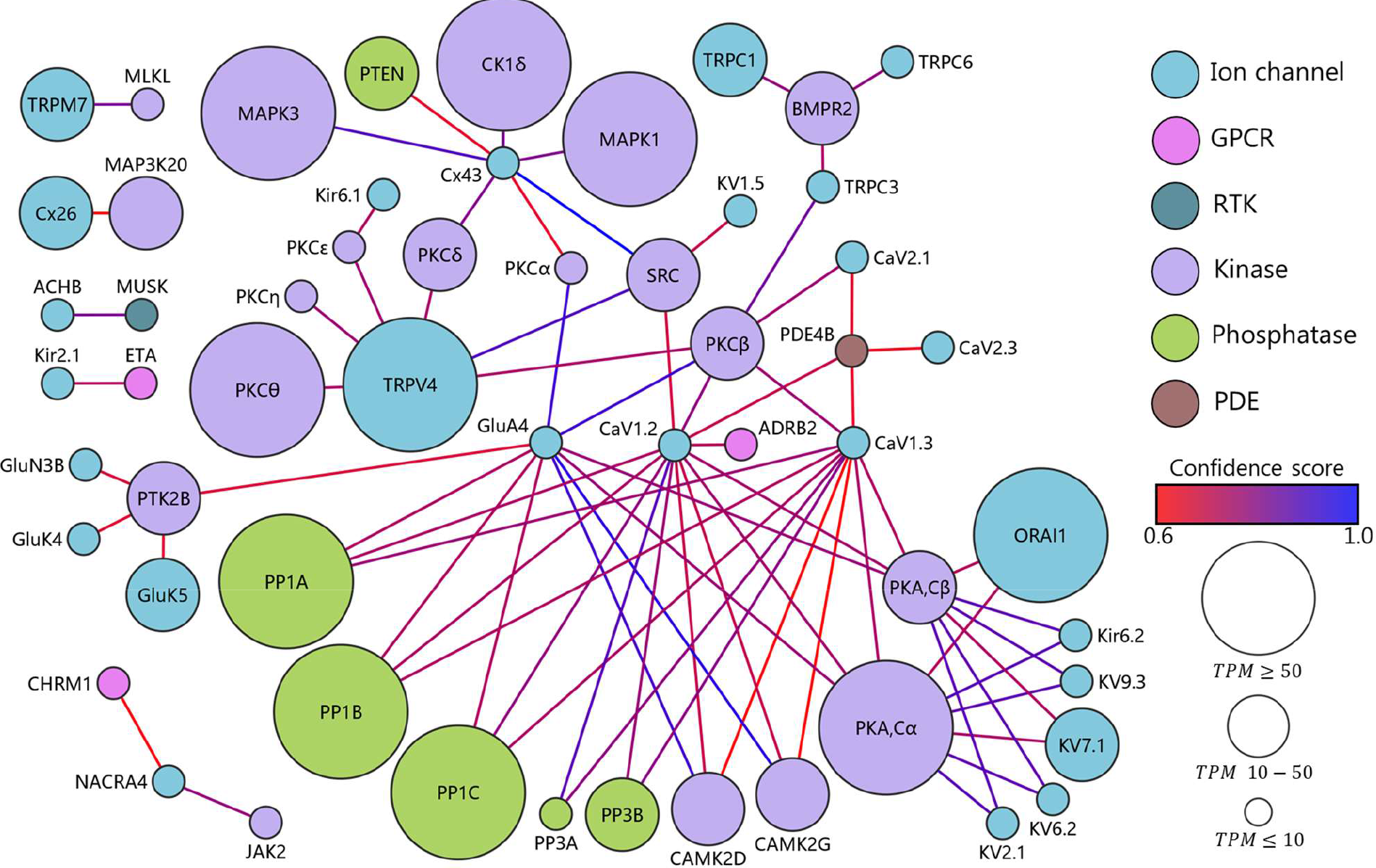
Network analysis of choroidal membrane channels and signaling cascades. Association network between plasma ion and water channels together with receptors (GPCRs and RTKs) and intracellular messengers (kinases, phosphatases, PDEs, and cyclases). The nodes (spheres) are colored based on their protein type. The size of the nodes corresponds to the transcriptional expression level in TPM and illustrates TPM ≤ 10, 10 -200, and ≥ 200. The confidence score is indicated by the color of the connecting lines from 0.7 (red) to 1.0 (blue). Only associations from empirical data and curated databases are included.

## Discussion

Here we reveal the choroidal abundance of transcripts encoding plasma membrane transport mechanisms that could potentially be involved in CSF secretion as well as those encoding regulatory factors that could partake in regulation of the CSF secretion machinery. The transcriptomic profile of male and female rat choroid plexus was nearly identical [65], as was the case for their transportome. The rat choroid plexus transcriptomic profile displayed high (∼ 90%) similarity to that of human and mice. The similarity may be even been higher than here reported, due to limited gene information of some of the retrieved genes.

Despite the fact that the existence of CSF and its continuous secretion from the blood to the brain have been long acknowledged, the exact molecular mechanisms by which this secretory process occurs have remained elusive [2, 12]. Several plasma membrane transport proteins are suggested to be implicated in the process, but their individual quantitative contribution remains unresolved. Several of these were detected amongst the 20 highest expressed membrane transport mechanisms revealed by filtering and curation of the RNAseq data obtained from rat choroid plexus; The Na^+^/K^+^- ATPase α1β1, NKCC1, TRPV4, AQP1, and the HCO_3_^-^ transporters NBCe2, NCBE, and AE2 [12, 42]. The transporter ranking was near-identical in RNAseq of entire choroid plexus and in choroid plexus epithelial cells captured by FACS, in support of the choroid plexus consisting predominantly of cells of epithelial origin [43, 44]. These ranked lists of choroidal transport mechanisms, in addition, provided gene names of other highly expressed transport proteins, which could potentially contribute to CSF secretion, but may never have been investigated for such a function. Of interest could be the highly expressed cation and anion transporters (BOCT, BSAT1), both of which are detected in brain barrier tissues [66, 67]. BOCT is expressed widely across many brain cell types, whereas the BSAT1 is expressed predominantly in choroid plexus and other barrier cell types, i.e. endothelial cells and pericytes [68]. The transported substrate of the former remains elusive [66], whereas the latter is involved in the transport of thyroxine, for which the choroid plexus is renowned [67]. The Na^+^-coupled sulfate transporter (SUT-1), ranked as number 6 highest expressed amongst transporters and pumps, is virtually exclusively expressed in choroid plexus amongst brain cells [69] (with some expression in the vascular leptomeningeal cells [68]). MCT8 and SNAT3 are both highly expressed in choroid plexus epithelial cells, but also detected in other brain cell types, albeit the latter predominantly in barrier- related cells, such as vascular endothelial cells, pericytes and ependymal cells [68]. MCT8 is involved in thyroxine transport [70] while SNAT3 contributes to the high glutamine content of the CSF [71]. None of these has, as of yet, been investigated for a potential implication in CSF secretion.

Comparison of the choroidal transcriptomic profile to that of another secretory epithelium of similar capacity, the kidney proximal tubules [2], revealed that choroid plexus expressed more than double the number of genes (∼ 13,500 genes) compared to the proximal tubules (∼ 6,000 genes, [72, 73]). 64% of the transport protein transcripts detected in the proximal tubules were also expressed in the choroid plexus, but these only amounted to 30% of those detected within this transcript category in the choroid plexus, suggesting that the choroid plexus serves a variety of tasks other than CSF secretion, some of which include transepithelial solute transport. Amongst the genes encoding transport proteins transcribed in both secretory epithelia, surprisingly few placed among highly expressed genes in both tissues: the Na^+^/K^+^-ATPase α1β1, AQP1, a voltage-gated Cl^-^ channel (ClC-4), in addition to two TRP channels (TRPML1 and TRPM7). Such similarity could suggest important roles of these particular transport proteins in the secretory processes (or regulation thereof) in these tissues [74–77], although, clearly, each epithelium appears to employ additional tissue-specific transport pathways to provide the transepithelial fluid transport. Of interest, an isoform of the Na^+^-coupled glucose transporter (SGLT2; SLC5A2), which was previously annotated to sole expression in the proximal tubules [78], was here detected in the choroidal transcriptome, albeit at a lower relative expression level (6 TPM) than that observed in the proximal tubule sample (22 TPM). The choroidal transcript abundance of SGLT2 is confirmatory of its recently demonstrated protein expression in mouse and human choroid plexus [79, 80]. The proposed selective SGLT2 expression in the proximal tubules led to development of SGLT2 inhibitors as a selective treatment option for type 2 diabetes mellitus [81, 82]. Such approach may have to be reconsidered based on SGLT2 expression in choroid plexus (this study and [79, 80]), where the transport protein could potentially partake in CSF secretion, like its homologue, SGLT1, participates in fluid transport across the small intestine [83].

The families of solute carriers (the SLCs) were highly represented in the transportome of the choroid plexus (approximately 63% of all the plasma membrane transport and pump protein transcripts), with most of the existing families expressed in this tissue (52 out of 66). Choroidal SLC expression is developmentally regulated, with notable upregulation of amino acid transporter families during embryonic stages [45, 84]. Grouping these SLC families into super categories defined by their transported substrate, we demonstrate that the electrolyte/HCO_3_^-^ and anion/cation transporters dominated at the transcript level. These were followed by amino acid, sugar, metal and vitamin transporters, in support of a role for choroid plexus in supplying the brain tissue with nutrients, micro-nutrients, and various co-factors [84, 85]. Notably, the choroid plexus is enriched in transcripts encoding metal transport proteins, with 22 genes dispersed among five different families of metal ion transporters. The choroidal expression of 15 different genes encoding various Zn^2+^ transporters of the efflux (SLC30; ZnT) and influx (SLC39, ZIP) types may serve to ensure transepithelial brain delivery of Zn^2+^ to various biochemical processes that are instrumental for proper brain development and function [84,86,87].

To reveal potential regulatory cascades involved in modulation of CSF secretion, we obtained lists of highly expressed plasma membrane receptors and signaling pathways expressed at the transcriptional level in choroid plexus. Amongst the GPCRs, the serotonin receptor of the 5-HT2C type was expressed at 4-fold higher abundance that the second-highest expressed receptor. The 5-HT2C receptor is expressed on the luminal side of the choroid plexus epithelium [88] and its activation leads to G_q_- dependent Ca^2+^ release from intracellular stores [89, 90], which subsequently promotes release of the insulin produced within the choroid plexus epithelium [90] and may modulate the rate of CSF secretion [91, 92]. The endothelin receptor B appears as the second highest expressed GPCRs, with subtype A further down the list, in support of their protein expression in the choroid plexus [93]. Endothelin may reduce the rate of CSF secretion [94], possibly via its action on the choroidal blood flow [95]. Of the receptor tyrosine kinases, growth factor receptors dominate the list of highest expressers, with three members of the family of fibroblast growth factor (FGF) receptors on the top 20 list (and two of these at the top). FGF receptors are detected at the protein level in choroid plexus [96] and FGFs may be implicated in brain fluid homeostasis by their ability to modulate NKCC1 activity [97] and to induce ventriculomegaly in a rodent model upon prolonged intraventricular infusion, at least in part due to formation of fibrosis and collagen deposits in the CSF drainage paths [98]. The latter observation aligns with the diminished foramen magnum area observed in hydrocephalic children bearing mutations in the gene encoding FGFR2 [99].

Lists of highly expressed intracellular signaling molecules include various cyclases, phosphodiesterases, kinases and phosphates. The vast majority of these remains to be associated with CSF secretion or regulation thereof, but may provide valuable hints to pursue in future efforts to modulate CSF secretion pharmacologically without targeting the choroidal transporters, many of which are expressed in other cell types or epithelia in the body. The cyclase-coupled receptors for atrial natriuretic peptide (ANPR-A and ANPR-B) were both detected amongst the top 10 highest expressed cyclases in the choroid plexus, and previously demonstrated at the protein level in this tissue [100]. ANP, via its induction of cGMP formation, may [100] or may not [101] cause decreased CSF secretion, and altered choroidal ANP receptor abundance in various forms of experimental hydrocephalus could indicate involvement in brain fluid dynamics [102]. Also of interest is the placement of the Ste20-related proline/alanine-rich kinase, SPAK, as the second highest expressed kinase in choroid plexus. SPAK is implicated in regulation of the CSF-secreting NKCC1 [6,62,63] and figures prominently in our network analysis as associated with NKCC1 as well as other cation, Cl^-^ cotransporters (CCCs). Other choroidal kinases and phosphatases associate with different transport mechanisms, i.e. various isoforms of the Na^+^/K^+^-ATPase and the TRPV4 channel, and may provide novel paths to investigate in future determinations of choroidal transport and its regulation.

As potential limitations to our study, we acknowledge the possibility that the lists of transporters, receptors, and intracellular signaling molecules are not absolute, as the information and annotation in the various databases, on which these are based, may be incomplete and with various levels of reliability. The filtration based on GO term annotation and manual curation was performed based on the available information and to the best of our knowledge. The network analysis is based on published work. Therefore, novel and unexplored connections between different transporters and their potential regulatory pathways are anticipated to be revealed by future research efforts. Lastly, transcript abundance may vary among the different choroid plexuses [25] and may not mirror the quantitative expression of proteins. Nevertheless, in the current study, we have created discovery tables of the transport mechanisms and regulatory pathways of the rat choroid plexus, and linked them via network analysis. We demonstrated high similarity between species (human and mouse) and sexes. The discovery tables provide lists of transport mechanisms that could participate in CSF secretion and suggest regulatory candidate genes that could be involved in their regulation. With these lists, we envision that researchers in the field may devise hypotheses regarding future quantification of transport mechanisms and their regulation, with the vision to obtain rational pharmaceutical targets for CSF production modulation in the pathologies involving disturbed brain water dynamics.

## Abbreviations

aCSF: Artificial cerebrospinal fluid
ATP: Adenosine triphosphate
cAMP: Cyclic adenosine monophosphate
CCC: Cation Cl- cotransporter
cGMP: Cyclic guanosine monophosphate
CSF: Cerebrospinal fluid
FACS: Fluorescence-activated cell sorting
FGF: Fibroblast growth factor
Genecards: Human gene database
GEO: Gene Expression Omnibus
GF: Growth factor
GO term: Gene Ontology term
GPCR: G protein-coupled receptor
HEPES: 4-(2-hydroxyethyl)-1-piperazineethanesulfonic acid
HGNC: Gene Nomenclature Committee
HUGO: Human Genome Organisation ICP Intracranial pressure
KEGG: Kyoto Encyclopedia of Genes and Genomes
NCBI: National Center for Biotechnology Information
PBS: Phosphate-buffered saline
PDE: Phosphodiesterase
RNAseq: RNA sequencing
RSEM: RNA-Seq by Expectation Maximization
RTK: Receptor tyrosine kinase
SLC: Solute carrier
STAR: Spliced Transcripts Alignment to a Reference
TPM: Transcripts per million
Uniprot: Universal protein resource

## Acknowledgements

We are grateful for the technical assistance from Trine Lind Devantier and Dagne Barbuskaite and for the valuable input on transcriptomics from Tune H Pers, Novo Nordic Center for Basic Metabolic Research, University of Copenhagen.

## Author contribution

SNA and NM designed the research study, SNA, TLTB, RV, and JHW carried out the experiments, SNA, TLTB, RV, JHW, and NM analyzed the data, SNA and NM drafted the manuscript and all authors approved the final version.

## Availability of data and materials

The datasets used and/or analyzed during the current study are available from the corresponding author on reasonable request.

Webserver database: https://cprnaseq.in.ku.dk

Scripts and data analysis: https://github.com/Sorennorge/MacAulayLab-RNAseq2

Raw data available at the NCBI GEO database with accession number: GSE194236 (https://www.ncbi.nlm.nih.gov/geo/query/acc.cgi?acc=GSE194236)

## Declarations

### Ethics approval

Animal experiments were in compliance with the European Community Council Directive 2010/63/EU on the Protection of Animals used for Scientific Purposes.

### Consent for publication

Not applicable

